# STRATEGIC IDENTIFICATION OF NEW GENETIC DIVERSITY TO EXPAND LENTIL *Lens culinaris* Medik. PRODUCTION USING NEPAL AS AN EXAMPLE

**DOI:** 10.1101/2020.08.18.256420

**Authors:** Sandesh Neupane, Rajeev Dhakal, Derek M. Wright, Deny Kumar Shrestha, Bishnu Dhakal, Kirstin E. Bett

## Abstract

This article is written to disseminate descriptive results obtained from field experiments conducted at Bardiya, Nepal, during winter 2016 and 2017 on 324 diverse lentil (*L. culinaris* Medik.) genotypes obtained from genebanks and breeding programs around the world. Lentil genotypes were examined for phenological traits related to adaptation, and recommendations for potential incorporation of new exotic genotypes into Nepalese breeding programs are presented. In addition, predictions of days to flowering based on temperature and photoperiod are used to identify genotypes that hold suitability for specific growing regions in Nepal, allowing for the potential expansion of growing regions. From this study we found many potentially adapted genotypes for terai, mid-hill and high hill growing regions, the list consists of a number of large-seeded lines with yellow cotyledons, an entirely new market class of lentils in Nepal. This paper primarily targets lentil breeders and agronomists; furthermore, it can be equally informative to extension workers involved in the pulse crop research and development in Nepal and other countries with similar climatic conditions.

## Introduction

Globally, lentil is cultivated in more than 50 countries (FAO 2020) typically categorized into three major lentil growing macro-environments: mediterranean, sub-tropical savannah, and temperate, where temperature and daylength differ considerably during the growing season (Tullu et al. 2011; Khazaei et al. 2016; Wright et al. 2020). There are about 58,000 accessions of the genus *Lens* (cultivated and wild species) currently housed in different gene banks worldwide (FAO 2010). These include landraces, breeding lines, advanced cultivars, and some unknown mixtures. Despite the availability of this large diversity, the majority of lentil breeding programs use only a fraction of it, primarily due to the adaptation constraints of lentil germplasm from one environment when grown in a different environment (Summerfield *et al*. 1985; Khazaei et al. 2016). These problems mostly arise from the temperature and photoperiod differences among environments (Summerfield et al. 1985). Therefore, in order to overcome adaptation constraints, understanding how diverse genotypes perform under field conditions is essential for sustainable lentil breeding. The systematic and judicious use of genetic variability helps maximize genetic gain and, over time, productivity.

Pulses are an integral part of a daily diet for Nepalese, and the majority of people consume pulses in the form of *Dal* at least twice a day. Lentil is ranked first in area and production amongst pulse crops, accounting for 62.93% of the total area and 65.76% of total pulse crop production in Nepal (MOALD 2020). It has also been prioritized as an agricultural commodity with high export potential, contributing to 2.3% of total national exports (USAID 2011). Although Nepal exports lentils to other parts of the world, the national production is not adequate to meet in-country demand and there is a large supply gap. With a few exceptions, lentil is produced throughout the country, however, commercial production is concentrated mostly in the terai (MOALD 2020), as the top ten lentil producing districts are in this region (Pokhrel et al. 2018). The twelve cultivars of lentil, predominantly small red types, which have been released through the formal system in Nepal (SQCC 2020) have been mostly grown by farmers in the terai region. However, they tend to yield far less in the farmers’ fields than global lentil yield would predict, suggesting room for improvement. There are numerous reasons for this gap including limited availability of quality seeds and technical knowledge, disease incidence (mostly Stemphylium blight and Fusarium wilt; Yadav et al. 2017), and climatic stresses, especially early and terminal drought and erratic rainfall (Magar et al. 2014). In addition, a dependency on limited genetic diversity (Ferguson et al. 1998; Khazaei et al. 2016), largely due to the genetic bottleneck created as lentils were disseminated through the Khyber Pass into the Indo-Gangetic Plain (Erskine et al. 2011), could be limiting increases in productivity. To overcome this constraint and expand germplasm diversity, the identification of germplasm sources that will likely be adapted to the region is necessary.

With few exceptions, recent research aimed at introducing new genotypes has focused on material obtained from ICARDA. Most of these have been of South Asia and Mediterranean origin, potentially due to the need for early flowering (Gharti et al. 2014; Yadav et al. 2016; Pokhrel et al. 2018). The inclusion of relatively early flowering exotic germplasm with higher yield potential from other regions might help to increase the current production and area. Embracing genotypes with more diverse seed size and cotyledon color may also create greater interest among farmers and consumers. Additionally, due to the broad range of lentil growing environments, there is room for expanding lentil production into new environments (e.g., high hills).

A photothermal model for lentil (1/f = a + bT + cP) described by Summerfield et al. (1985), has the potential for predicting days from sowing to flowering (DTF) and has been used in a wide range of pulse crops. A recent study by Wright et al. (2020) applied this photothermal model to a diverse collection of 324 lentil genotypes, enabling prediction of DTF of a given genotype by providing the temperature and photoperiod information of a chosen location. In addition, observations of DTF across nine locations were used to cluster genotypes into eight groups based on their phenological response to differing environments. The aim of this study was to identify sources of new diversity for lentil research and cultivar development in Nepal. With the limited availability of a narrow range of local as well as introduced diversity, the identification of promising, additional adapted genotypes can be employed with immediate action to aid in cultivar choice among farmers and potentially expand the area for lentil production in Nepal. More importantly, the information generated from this study could be used in breeding programmes with a view to future cultivar development through crosses with novel sources of diversity.

## Materials and methods

### Field experiments and data collection

Two field trials were conducted in the winter of 2016 and 2017 at Bardiya, Nepal (28°15’07.6”N, 81°30’05.4”E) with a diversity panel of 324 lentil genotypes obtained from genebanks of the International Center for Agricultural Research in the Dry Areas (ICARDA), United States Department of Agriculture (USDA), Plant Gene Resources of Canada (PGRC), as well as cultivars developed at the Crop Development Centre (CDC) of the University of Saskatchewan (U of S) (*https://knowpulse.usask.ca/Lentil-Diversity-Panel)*. These genotypes primarily originated from 43 different countries, along with a few breeding lines from ICARDA, representing all three major lentil growing environments. The panel varied in seed size, cotyledon and seed coat colors and phenological traits. Seeds of each genotype were sown in small plots with 25 seeds in two 1 m rows in the first year and 50 seeds in two 1 m rows (50 cm apart) in the second year. For both years, the experimental design was a randomized lattice square (18 × 18), replicated three times. Both field experiments were phenotyped for Days to - emergence (DTE), flowering (DTF), swollen pod (DTS) and maturity (DTM) when 10% of plants had emerged, flowered, had swollen pods, and had 50% dry pods, respectively. Vegetative period (VEG) was recorded as the number of days between emergence and flowering and the reproductive period (REP) was calculated as the difference between DTM and DTF. Environmental data, temperature and rainfall, were obtained from the meteorological station located at Khajura, Banke (28°6’35”N, 81°35’42”E) in both years. Photoperiod data were extracted using the ‘insol’ package in R (Corripio 2019) after providing latitude, longitude, specific day and time zone; duration between sunrise to sunset was used as the photoperiod value.

### Data analysis

All data analyses were performed in R 3.5.0 software (R Core Team 2018). Data wrangling and visualization was done using the R packages: ‘tidyverse’ (Wickham 2017), ‘skimr’ (Quinn et al. 2019), ‘ggbeeswarm’ (Clarke and Sherrill-Mix 2017), ‘ggpubr’ (Kassambara 2018). An analysis of variance (ANOVA) was performed with the mixed model procedure using the ‘lmerTest’ package (Kuznetsova et al. 2017) to compare 324 lentil genotypes for DTE, DTF, DTS, DTM, VEG and REP. During the analysis, genotype (G), experiment year (E) and their interaction (G X E) were considered as the fixed effects, whereas block nested over the replication again nested within the experiment year was considered as a random effect. The mixed model we used for the analysis was -

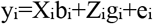

where y_i_ is a response variable observed for individual i, b_i_ is a vector of fixed effects, g_i_ is a vector of random effect for individuals and e_i_ represents residuals for the trait or environment i. X and Z are explanatory variables.

For each phenological trait, if the ANOVA indicated significant differences at or above the *P<0.05* level, the means were separated by the least significant difference (LSD) method using “emmeans’ package (Russell 2019) in R. Values of measured traits were averaged across the two years if no significant G X E (*P>0.05*) was detected. For analysis of individual years where G X E was significant, genotypes were considered a fixed effect, and replication was considered a random effect.

Correlation analysis among phenological traits was conducted using Pearson’s method and plots were generated using the ‘GGally’ (Schloerke et al. 2019) package in R.

## Results and discussion

### Variation in phenological traits among genotypes and experimental years

Considerable variation was observed for phenological traits among 324 lentil genotypes as well as between years (Fig. 1). The phenological traits DTE, DTF, DTS and DTM were all significantly different (*P<0.001*) for genotype and had a significant genotype by year interaction (Table 1). The vegetative period (VEG) and reproductive period (REP) were also significantly different (*P<0.001*) for genotype and had a genotype by year interaction. All traits, except DTE and REP, were significantly different between years. The significant year variation for DTF, DTS and DTM are likely the result of environmental factors related to an earlier seeding date in the second year (Fig. 2). As both field experiments were conducted during the same time of year and at the same location, photoperiod did not differ significantly (Fig 2); thus, the phenotypic variation was most likely due to the temperature, rainfall and other additional factors such as light quality, solar radiation, etc. experienced in the second growing year. Due to elevated temperatures late in the growing season, 67 genotypes did not flower, and an additional 153 genotypes did not make it to full maturity in at least one replication in 2016. At 149 days after sowing, all plots that produced seeds were harvested in 2016.

**Figure 1.**
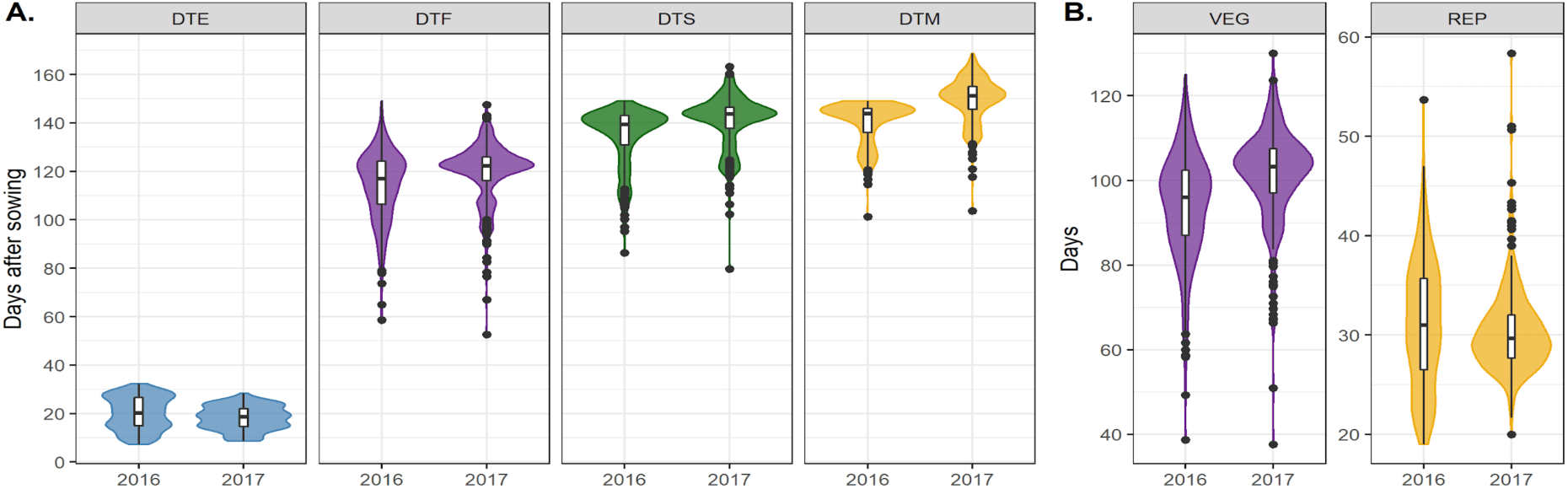
Variation in **A.** Days to – emergence (DTE), flowering (DTF), swollen pod (DTS), maturity (DTM) from seeding, and **B.** vegetative period (VEG) and reproductive period (REP) of 324 genotypes grown in the winter of 2016 and 2017 at Bardiya, Nepal. The width of a plot indicates the distributions’ density, and the whiskers on the box plots represent 1.5 times the quartile of the data. Individuals falling outside the range of the whiskers are represented as black dots.

**Figure 2.**
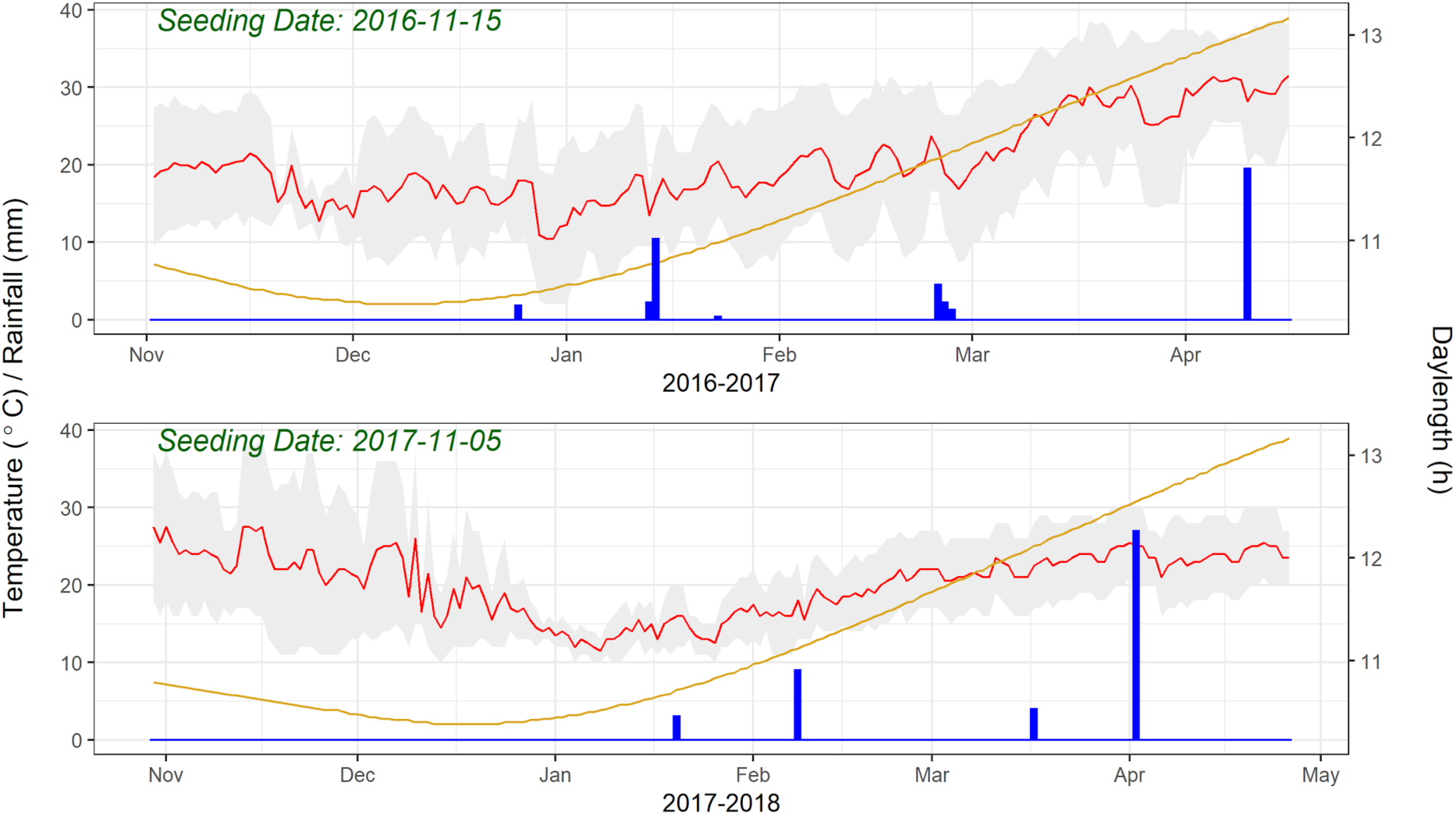
Variation in temperature (°C), rainfall (mm) and daylength (h) during field experiments in 2016 (top) and 2017 (bottom). The X-axis represents the months of the field experiments, and Y-axis represents temperature, rainfall (precipitation) and daylength. The red line in both plots is the average temperature, and the grey shadow on the red line is maximum and minimum temperature, whereas the yellow line is daylength, and blue bars are the amount of rainfall on a particular day.

**Table 1.** Analysis of Variance (ANOVA) results of 324 lentil genotypes evaluated for DTE, DTF, DTS and DTM as well as VEG and REP in the winter of 2016 and 2017 at Bardiya, Nepal.

| Source of Variation | DTE | DTF | DTS | DTM | VEG | REP |
| --- | --- | --- | --- | --- | --- | --- |
| Genotype | *** | *** | *** | *** | *** | *** |
| Year | ns | * | ** | *** | ** | ns |
| Genotype X Year (G X E) | *** | *** | *** | *** | *** | *** |

DTE=Days to Emergence, DTF=Days to Flowering, DTS=Days to Swollen Pod, DTM=Days to Maturity, VEG=Vegetative Period and REP= Reproductive Period
\*\*\*, significant at $P < 0.001$ ; \*\*, significant at $P < 0.01$ ; \*, significant at $P < 0.05$ ; ns, not significant.

The strong positive correlations of DTF with DTS, DTM and VEG, and negative correlation with REP (Fig. 3), as well as a non-signifícant difference of REP among years (Table 1), indicated DTF is the primary factor driving adaptation and selection of genotypes in a Nepalese environment. Therefore, only DTF is used hereafter for further discussion.

**Figure 3.**
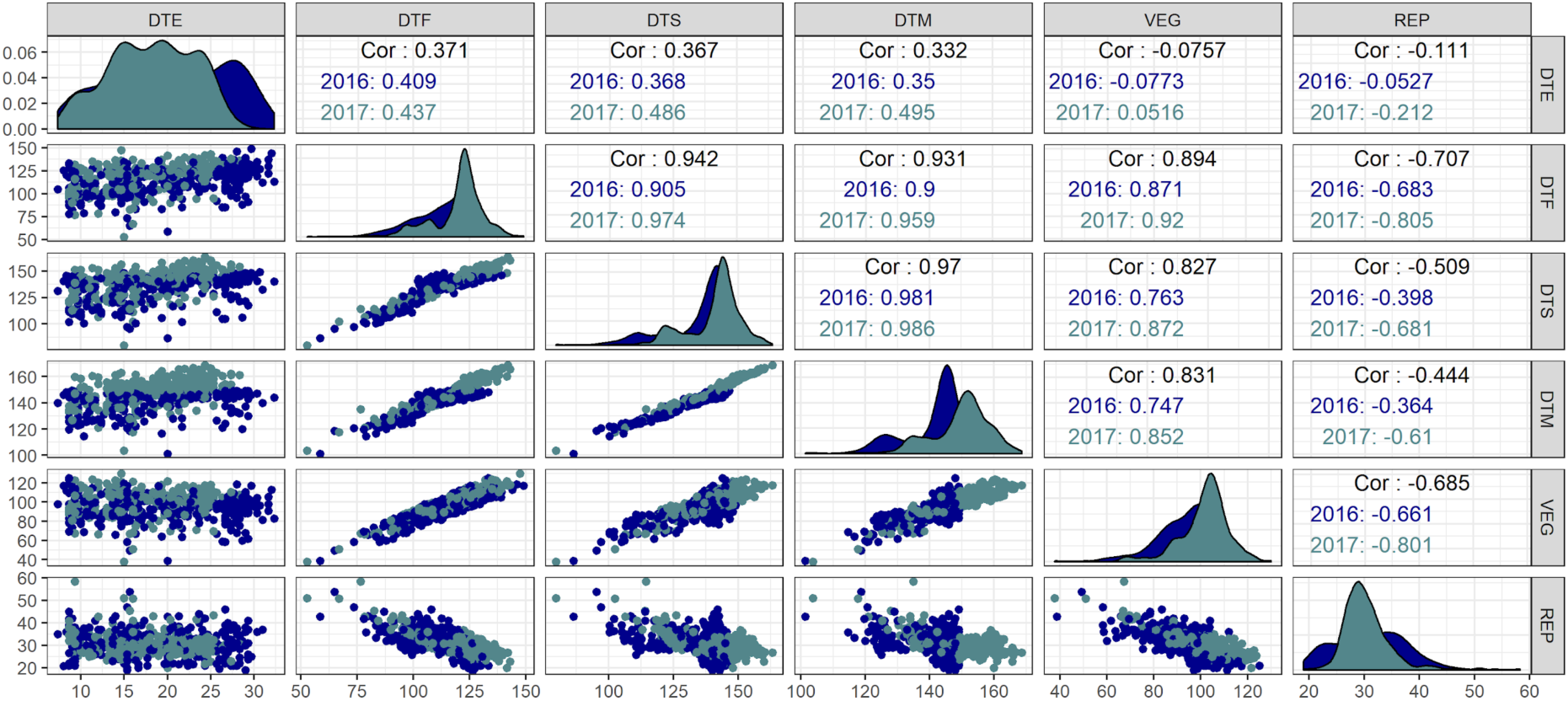
Correlations along with the corresponding correlation coefficients (shown as Cor) among days from sowing to - emergence (DTE), flowering (DTF), swollen pod (DTS), maturity (DTM) as well as vegetative period (VEG) and reproductive period (REP). Different colors represent two different years of field trials, i.e. 2016 (dark blue) and 2017 (grey-blue).

### New genotypic options for Nepalese agricultural systems

Variations in DTF existed both within and amongst countries of origin, with genotypes from South Asian countries tending to flower earliest, those from Latin American and temperate countries the latest, and both early and later flowering genotypes came from Mediterranean countries (Fig. 4). However, caution should be used when considering the country of origin, as this information might not always be correct and could represent diverse growing regions that exist within a country. For example, Ethiopia, Turkey, and Nepal have both highland and lowland production regions with differing temperature and photoperiod regimes during the growing season.

**Figure 4.**
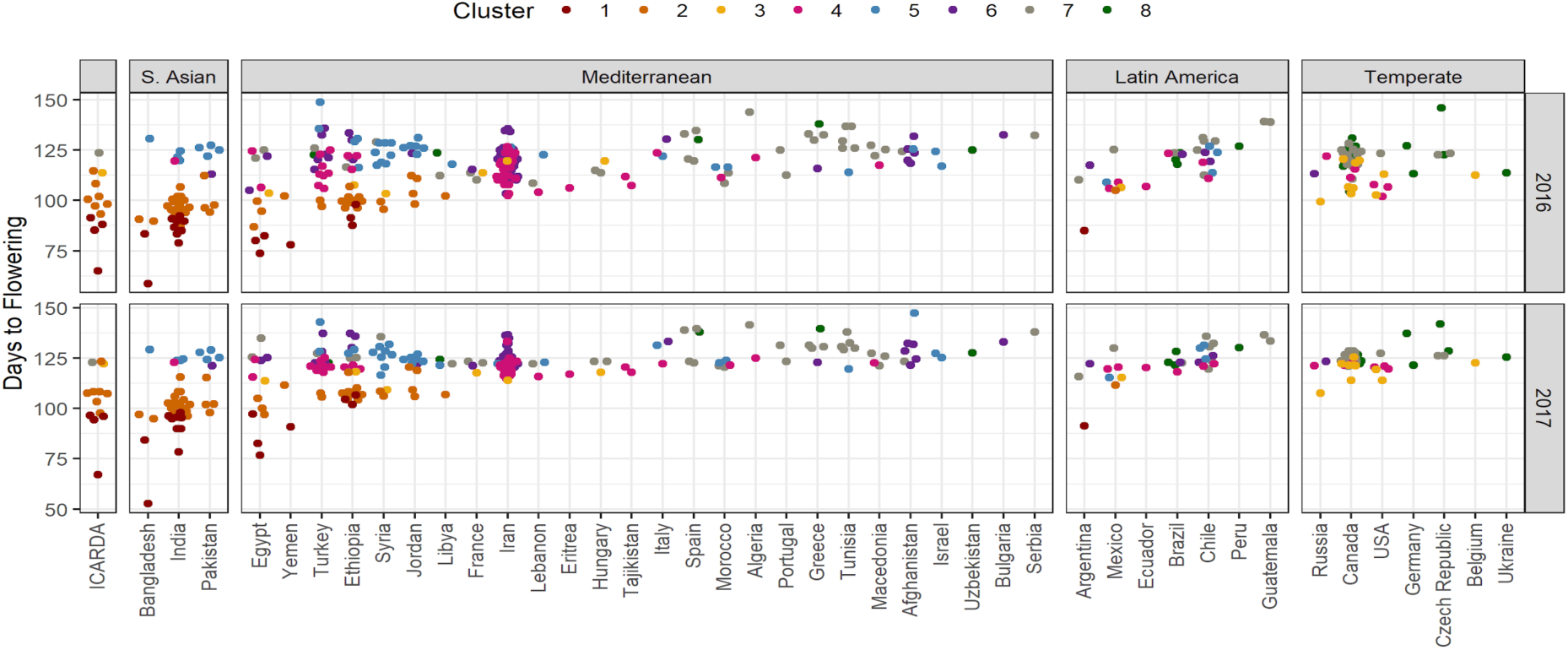
Days to Flowering (DTF) of 324 genotypes grown in the winter of 2016 and 2017 at Bardiya, Nepal, based on their country and geographical region of origin. Different colored dots represent genotypes from eight different clusters derived from principal component analysis and hierarchical clustering (Wright et al. 2020). Genotypes from the ICARDA breeding program were kept separate as they are a mix of germplasm from a large breeding program.

Using principal component analysis (PCA) and hierarchical clustering of DTF across 18 site-years, the 324 genotypes were classified into eight groups, reflecting their phenological response to varying temperatures and photoperiods (Wright et al. 2020). Genotypes with South Asian origins were predominately found in clusters 1 and 2, with some later flowering genotypes in cluster 5 (Fig. 4). Cluster 5 represents those genotypes with the lowest temperature sensitivity (Wright et al. 2020), likely adapted to the cooler, intermediate elevations between the Afghanistan highlands and the Indo-Gangetic Plain as lentils were disseminated through the Khyber Pass. The lack of representation of south Asian genotypes in other clusters illustrates the intense selection pressures that have resulted in a bottleneck limiting genetic diversity within germplasm grown in the region. These clusters can be used as a guide for identifying new genotypes that could be used to increase production and genetic diversity in the region.

Among the eight DTF clusters, cluster 1 was earliest to flower followed by cluster 2, then cluster 3; clusters 4-8 were all similar and late (Fig. 4). Due to the need of lentil breeders in South Asia to develop early flowering and maturing varieties to capture residual soil moisture and to avoid spring terminal drought (Sarker and Erskine 2006), all genotypes from clusters 1 and 2 could be considered for further screening and immediate use in a breeding program. If, on the other hand, the goal is to broaden the genetic diversity, identifying genotypes outside of these cluster groups would be preferred but would need to be crossed to adapted material and subject to intense selection for appropriate maturity. Likewise, since a majority of the lentil cultivars in Nepal originated from either South Asia or the Mediterranean (Gharti et al. 2014; Yadav et al. 2016; Pokharel et al. 2018), genotypes originating from beyond the usual sources (i.e. countries other than South Asian, Mediterranean, and ICARDA origin), but that still flower and mature early, would also serve this purpose.

### Identifying genotypes for testing in expanded growing regions using a photothermal model

In a previous study by Wright et al. (2020), a photothermal model was used to predict DTF on the 324 lentil genotypes used in this study based on multiple field experiments in the major lentil growing regions. This knowledge can be leveraged to predict DTF based on the average temperature and photoperiod of a given environment and identify genotypes which likely have the appropriate phenology for that location. Since the Grain Legume Research Program of Nepal is located at Khajura, Banke, and the Nepal Agricultural Research Council has research stations located in Kathmandu and Jumla where they have been conducting lentil research trials, these three locations were chosen as representative sites of the major agricultural growing regions - the terai, mid hills and high hills, respectively. The average temperature during the lentil growing season in these three locations (from 1982 to 2012) was acquired from https://en.climate-data.org/. We then ran the photothermal model to identify putative genotypes suitable for testing in these environmentally contrasting growing regions. Due to the tendency for under-prediction of DTF by the model in Nepal (Wright et al. 2020), the desire for early flowering by breeders, and the dominance of clusters 1, 2 and 5 among South Asian genotypes, we used the maximum DTF from cluster 2 of the South Asian genotypes as a cut off for our selection criteria when evaluating for potentially adapted genotypes in Banke and Kathmandu. Similarly, we used the maximum DTF of cluster 5 for Jumla since we predict those likely represent the highland adapted genotypes.

In Banke, lentil is generally seeded in early November and harvested around late March to early April (Yadav et al. 2007). The average temperature during the lentil growing season would be around 20.1°C, and the day length is 11.16 h. After entering this information into the model, we found 99 genotypes (Fig. 5), which might be suitable for testing in the terai region. 71 of these are from outside South Asia; 32 are not from clusters 1 or 2, and 14 are large-seeded with yellow cotyledons offering a range of diversity (Supplemental Table 1). Similar to Banke, lentil is generally seeded around late October to early November and harvested around late March to early April in and around the Kathmandu Valley (Neupane and Shrestha 1991). The average temperature during this time would be around 13.8 °C and daylength is 11.2 h. Based on the model, we found 82 genotypes (Fig. 5) which might be suitable for the mid-hills region. 51 of these are originating from outside South Asia; 10 are not from clusters 1 or 2, and one is large-seeded with yellow cotyledons (Supplemental Table 2). Unlike in Banke and Kathmandu, lentil is generally seeded late March to early April and harvested in late August to early September in Jumla (Dhital et al. 1994). The average temperature during this time would be 9.6 °C with an average daylength of 13.5 h. Based on the model, we found 126 genotypes (Fig. 5) which might be suitable for the high hills region. 91 of these originated from outside South Asia; 56 of these are not from clusters 1 or 2, and 34 are large-seeded with yellow cotyledons mostly from cluster 5, a group with reduced sensitivity to low temperatures (Supplemental Table 3).

**Figure 5:**
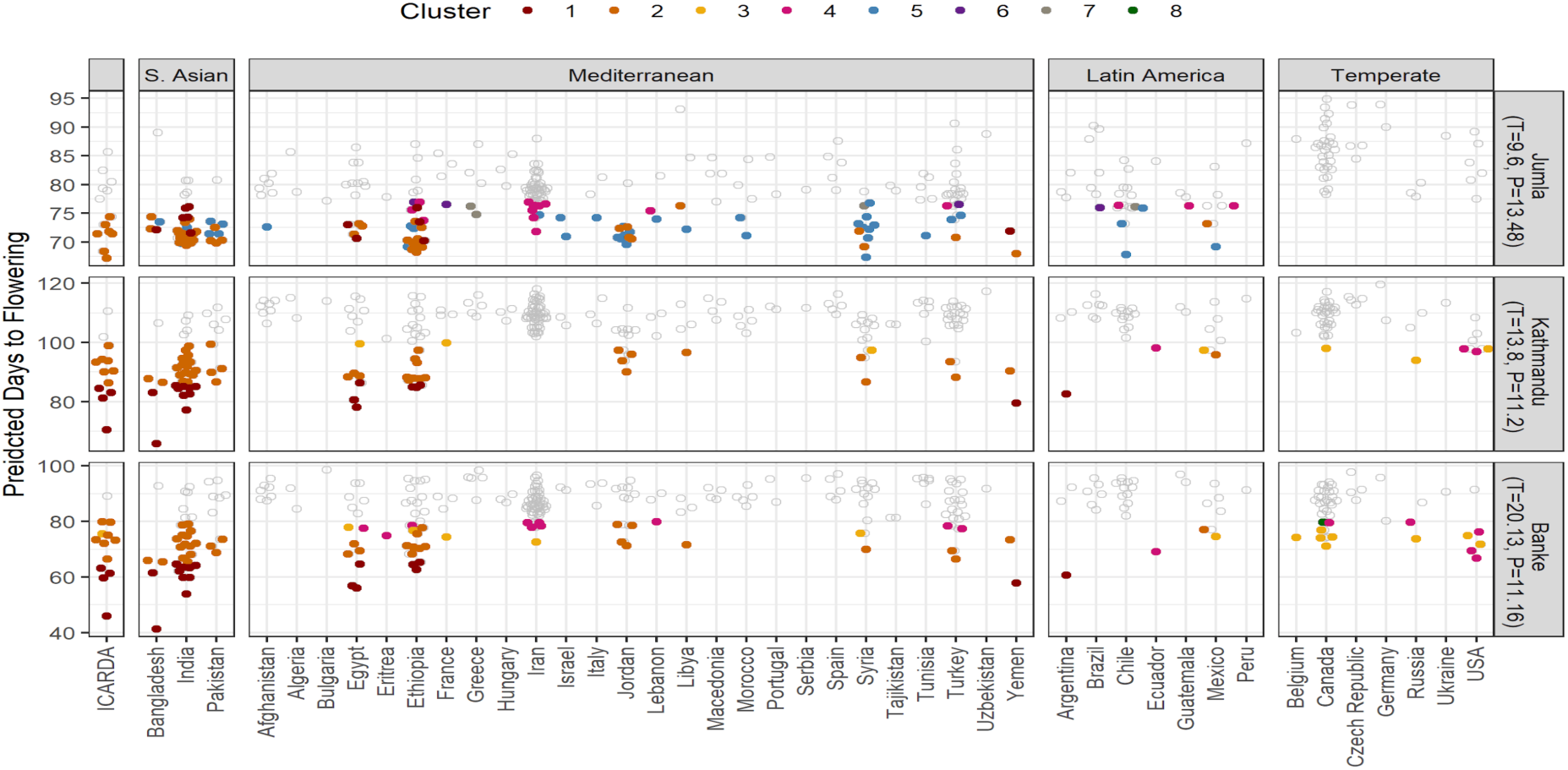
Predicted Days to Flowering (DTF) of 324 genotypes for Banke, Kathmandu, and Jumla as the representative of the major agricultural growing regions viz. the terai, mid-hill and high hill of Nepal, derived from a photothermal model from the study by Wright et al. (2020) using in-season average temperature (T) and photoperiod (P) of each environment. The colored points are the genotypes from different clusters, considered to have appropriate phenology for the location or adapted genotypes out of 324 genotypes set for each growing region. For prediction, the maximum DTF from cluster 2 of the South Asian genotypes was considered a cutoff for Banke and Kathmandu, the maximum from cluster 5 for Jumla.

These genotypes should be tested on-farm and could be used to establish rapid expansion of the lentil production in high hill regions. In regions like Jumla where lentils would be grown in spring to fall, it had been expected that genotypes from temperate regions (e.g., Canada) might perform better than the genotypes from winter growing regions (e.g. India); however, we did not find such results from the photothermal model (Fig. 5). This emphasizes the strong role of the combination of temperature and photoperiod in lentil adaptation and demonstrates the potential usefulness of the photothermal model for pre-screening genotypes.

Among the adapted genotypes from different clusters, genotypes from clusters 1 and 2 were deemed more promising than those from other groups for both highland and lowland locations. Cluster 5 offers additional genotypes that may be suitable in the highland location, supporting the idea that the highlands were an intermediate production region for lentil as they were disseminated into South Asia, and were selected for early flowering (Erskine et al. 2011). In all three locations, we have identified a set of adapted genotypes within which are genotypes with larger seed sizes and different cotyledon colors than the currently available Nepalese cultivars. This would help to establish additional market classes for export from Nepal.

While this analysis focused on DTF, and thus does not directly address the yield gap present between Nepal and some of the other major production regions for lentils, the introduction of new, exotic genetic diversity, identified herein should assist and/or increase the yearly yield gain by lentil breeders in Nepal. We recommend separate trials with these genotypes in larger sized plots to assess the grain yield.

## Conclusion

Significant variation for all phenology related traits was present among the 324 lentil genotypes grown in Bardiya, Nepal. Days to flower was considered the primary factor governing adaptation due to a high positive correlation with maturity, and a negative correlation with the reproductive period. A previous study involving this diversity panel categorized lentil genotypes into eight groups and demonstrated the ability to predict DTF based on average temperature and photoperiod. Leveraging this knowledge, we were able to predict genotypes suitable for different lentil production regions within Nepal that could be used to expand the genetic diversity. When considering Banke, Kathmandu and Jumla as the representative of terai, mid-hill and high hill growing regions, we found many potentially adapted genotypes for each region and several of which are either large-seeded or with yellow cotyledons that could contribute to entirely new market classes of lentils in Nepal and help increase production and export options.

## Declarations

## Authors’ contributions

K.E.B., S.N. and D.W. conceived and designed the study. D.K.S., B.D., R.D. and S.N. managed the field trials and collected phenotypic data. S.N. analyzed phenotypic data. S.N, D.W., R.D. and K.E.B. wrote the manuscript. All authors reviewed and approved the manuscript.

## Acknowledgement

We are grateful to all funding agencies and collaborative partners of project AGILE. Our special thanks to Ms. Crystal Chan for managing the project and supporting us in every step throughout the field experiments, and beyond. We acknowledge the support from Local Initiatives for Biodiversity, Research and Development (LI-BIRD) for conducting the field experiments. Thanks to Mr. Drona Kumar Shrestha, Ms. Phool Kumari Rai, Ms. Samjhana Magar and Mr. Baikuntha Adhikari for their support in managing day to day work at the field. We are thankful to Dr. Rajendra Darai and Mr. Laxman Aryal from the Grain Legumes Research Program (GLRP) under the Nepal Agricultural Research Council (NARC) for technical support during the field experiments.

## Supplemental tables

Table 1: Germplasm listing along with phenological data collected over two years of testing in Bardiya, Nepal as well as seed characteristics.

Table 2: Environmental data – daily max and min temperatures, mean daily temperature, rainfall amounts and daylength during the growing seasons over two years of testing in Bardiya, Nepal.

Table 3: List of potential genotypes for Banke, Nepal, representative of terai growing region, obtained after providing in-season average temperature and photoperiod, expected days to flowering, and using the coefficients derived phenological study in lentil by Wright et al. (2020) into the photothermal model 1/f = a +bT +cP.

Table 4: List of potential genotypes for Kathmandu, Nepal, representative of mid-hill growing region, obtained after providing in-season average temperature and photoperiod, expected days to flowering, and using the coefficients derived phenological study in lentil by Wright et al. (2020) into the photothermal model 1/f = a +bT +cP.

Table 3: List of potential genotypes for Jumla, Nepal, representative of high hill growing region, obtained after providing in-season average temperature and photoperiod, expected days to flowering, and using the coefficients derived phenological study in lentil by Wright et al. (2020) into the photothermal model 1/f = a +bT +cP.

## References

Clarke E, Sherrill-Mix S (2017) Ggbeeswarm: Categorical Scatter (Violin Point) Plots. https://CRAN.R-project.org/package=ggbeeswarm

Corripio JG (2019) Insol: Solar Radiation. https://CRAN.R-project.org/package=insol

Dhital BK, Budhathoki CB, Devkota HP (1994) Time to planting trails of cowpea and lentil. Lumle Agricultural Research Centre, Pokhara, Nepal

Erskine W, Sarker A, Ashraf M (2011) Reconstructing an ancient bottleneck of the movement of the lentil (*Lens culinaris* ssp. culinaris) into South Asia. Genetic Resources and Crop Evolution. 58: 373–381

FAO (2010) The State of the World’s Plant Genetic Resources for Food and Agriculture. FAO, Rome. http://www.fao.org/docrep/013/i1500e/i1500e00.htm. Accessed 18 March 2020

FAO (2020) FAO Statistics Database. http://www.fao.org/faostat/en/#data/QC/visualize Accessed 14 Feb 2020

Ferguson ME, Robertson LD, Ford-Lloyd BV, Newbury HJ, Maxted N (1998) Contrasting genetic variation amongst lentil landraces from different geographical origins. Euphytica. 102, 265–273. https://doi.org/10.1023/A:1018331432580

Gharti DB, Darai R. Subedi S, Sarker A, Kumar S (2014) Grain Legumes in Nepal: Present Scenario and Future Prospects. World Journal of Agricultural Research. 2(5): 216–222

Kassambara A (2018) ggpubr: ‘ggplot2’ Based Publication Ready Plots. https://CRAN.R-project.org/package=ggpubr

Khazaei H, Caron CT, Fedoruk M, Diapari M, Vandenberg A, Coyne CJ, McGee R, Bett KE (2016) Genetic Diversity of Cultivated Lentil (Lens culinaris Medik.) and Its Relation to the World’s Agro-ecological Zones. Frontiers in Plant Science. 7: 1093

Kuznetsova A, Brockhoff PB, Christensen RHB (2017). lmerTest Package: Tests in Linear Mixed Effects Models. Journal of Statistical Software. 82(13): 1–26

Magar DBT, Darai R, Gauchan D, Sarker A (2014) Varietal adoption and marketing of lentil in the mid and far western terai region of Nepal. Adv Plants Agric Res. 1(5): 164–170

MOALD (2020) Statistical Information on Nepalese Agriculture 2075/76 [2018/19]. Ministry of Agriculture and Livestock Development, Singhadurbar, Kathmandu, Nepal. https://www.moald.gov.np/publication/Agriculture%20Statistics. Accessed 10 August 2020

Neupane RK, Shrestha R, (1991) Report on lentil varietal and agronomical research. Nepal Agricultural Research Council, Lalitpur, Nepal

Pokhrel A, Aryal L, Poudel PP (2018). A review on research work of grain legumes research program, NARC. Government of Nepal, Nepal Agricultural Research Council, Grain Legume Research Program, Banke, Nepal

Quinn M, McNamara A, de la Rubia EA, Zhu H, Ellis H (2019) skimr: Compact and Flexible Summaries of Data. R package version 1.0.7. https://CRAN.R-project.org/package=skimr

R Core Team. (2018) R: A language and environment for statistical computing. R Foundation for Statistical Computing, Vienna, Austria. https://www.R-project.org/

Russell L (2019) emmeans: Estimated Marginal Means, aka Least-Squares Means. R package version 1.3.4. https://CRAN.R-project.org/package=emmeans

Sarker A, Erskine W (2006) Recent Progress in the ancient lentil. Journal of Agricultural Science. 144: 19–29

Schloerke B, Crowley J, Cook D, Briatte F, Marbach M, Thoen E, Elberg A, Larmarange J (2019) GGally: Extension to ‘Ggplot2’. https://github.com/ggobi/ggally

SQCC (2020) Notified and Denotified Varieties till 2076 04 07. Seed Quality Control Centre, Ministry of Agriculture and Livestock Development, Singhadurbar, Kathmandu, Nepal. http://sqcc.gov.np/images/category/Notified_and_Denotified_Varieties_till_2076_04_07.pdf

Summerfield RJ, Roberts EH, Erskine W, Ellis RH (1985) Effects of temperature and photoperiod on flowering in lentils (Lens culinaris Medic.). Annals of Botany. 56(5): 659–671

Tullu A, Diederichsen A, Suvorova G, Vandenberg A (2011) Genetic and genomic resources of lentil: status, use and prospects. Plant Genet. Resour. 9: 19–29

USAID (2011) Nepal Economic Agriculture, and Trade Activity Value Chain/ Market Analysis of the Lentil Sub-Sector in Nepal. United States Agency for International Development, General Development Office, Kathmandu, Nepal.

Wickham H (2017) Tidyverse: Easily Install and Load the ‘Tidyverse’. https://CRAN.R-project.org/package=tidyverse.

Wright DM, Neupane S et al (2020) Understanding photothermal interactions will help expand production range and increase genetic diversity of lentil (*Lens culinaris* Medik.). bioRxiv. https://doi.org/10.1101/2020.07.18.207761

Yadav SS, Rozvi AH, Manohar M, Verma AK, Shrestha R, Chen C, Bejiga G, Chen W, Yadav M, Bahl PN. In. SS Yadav et al. (eds.) 2007. Lentil: An Ancient Crop for Modern Times. Springer. 415–442.

Yadav N, Ghimire S, Shrestha S, Sah B, Sarker A, Sah S (2017) Source of resistant against Fusarium wilt and Stemphylium blight in lentil (*Lens culinaris* Medikus). International Journal of Applied Sciences and Biotechnology. 5(1), 102–107.

